# Advancing functional genetics through *Agrobacterium*-mediated insertional mutagenesis and CRISPR/Cas9 in the commensal and pathogenic yeast *Malassezia furfur*

**DOI:** 10.1101/638429

**Authors:** Giuseppe Ianiri, Gabriel Dagotto, Joseph Heitman

**Affiliations:** Department of Molecular Genetics and Microbiology, Duke University Medical Center, Durham, NC, 27710, USA

**Keywords:** *Malassezia furfur*, insertional mutagenesis, CRISPR/Cas9, protein phosphatase 2A, pleiotropic drug resistance

## Abstract

*Malassezia* encompasses a monophyletic group of basidiomycetous yeasts naturally found on the skin of humans and other animals. *Malassezia* species have lost genes for lipid biosynthesis, and are therefore lipid-dependent and difficult to manipulate under laboratory conditions. In this study we applied a recently-developed *Agrobacterium tumefaciens*-mediated transformation protocol to perform T-DNA random insertional mutagenesis in *Malassezia furfur*. A total of 767 transformants were screened after exposure to 10 different stresses, and the 19 mutants that exhibited a phenotype different from the wild type were further characterized. The majority of these strains had single T-DNA insertions, which were identified within the open reading frames of genes, within untranslated regions, and in intergenic regions. Some T-DNA insertions generated chromosomal rearrangements, and others could not be characterized. To validate the findings of the forward genetic screen, a novel CRISPR/Cas9 system was developed to generate targeted deletion mutants for 2 genes identified in the screen: *CDC55* and *PDR10*. This system is based on co-transformation of *M. furfur* mediated by *A. tumefaciens* to deliver both a *CAS9*-gRNA construct that induces double-strand DNA breaks, and a gene replacement allele that serves as a homology directed repair template. Targeted deletion mutants for both *CDC55* and *PDR10* were readily generated with this method. This study demonstrates the feasibility and reliability of *A. tumefaciens*-mediated transformation to aid in the identification of gene functions in *M. furfur* through both insertional mutagenesis and CRISPR/Cas9-mediated targeted gene deletion.

## Introduction

The genus *Malassezia* is a lipophilic, monophyletic group of basidiomycetous yeasts that colonize sebaceous skin sites and represents more than 90% of the skin mycobiome (Findley *et al*. 2013; Wu *et al*. 2015; Byrd *et al*. 2018). In addition to a ubiquitous presence on the skin of human and animals, recent data support the hypothesis that *Malassezia* fungi are much more widespread than previously thought. Metagenomics studies revealed the presence of *Malassezia* DNA in a number of unexpected areas such as in association with corals and sea sponges in the ocean, although *Malassezia* marine species have yet to be isolated in axenic culture (Amend *et al*. 2019). There are currently 18 species within the *Malassezia* genus. One defining characteristic of the *Malassezia* genus is the lack of a fatty acid synthase, Δ^9^ desaturase, and Δ^2,3^ enoyl CoA isomerase, making them lipid-dependent and difficult to study and manipulate under laboratory conditions. *Malassezia* are highly divergent from other fungi that are commonly found on the skin, such as *Candida* species and the dermatophytes. Furthermore, *Malassezia* species belong to the Ustilaginomycotina subphylum, which includes the plant pathogens *Ustilago, Sporisorium*, and *Tilletia*, and are highly divergent from other basidiomycetous fungi that infect humans, such as *Cryptococcus neoformans*. Recent classifications revealed that *Malassezia* represents a sister group to the blast yeast-like fungi *Moniliella* (Wang *et al*. 2014; Wang *et al*. 2015), which includes species reported to be pathogenic on human and animal skin (McKenzie *et al*. 1984; Pawar *et al*. 2002) as well as others that are of interest in sugar alcohol production in industrial settings (Kobayashi *et al*. 2015).

In the last decade, there has been increasing scientific interest in *Malassezia*, with several sequencing projects aimed at defining genomic features and gene content for 15 broadly recognized *Malassezia* species (Xu *et al*. 2007; Gioti *et al*. 2013; Triana *et al*. 2015; Wu *et al*. 2015; Park *et al*. 2017; Zhu *et al*. 2017; Kim *et al*. 2018; Lorch *et al*. 2018; Cho *et al*. 2019; Morand *et al*. 2019). All haploid *Malassezia* species have small genomes compared to other phylogenetically related fungi (7 to 9 Mb compared to ~20 Mb), and have lost genes involved in carbohydrate metabolic processes and hydrolysis activity. Genome analyses have revealed intriguing features, such as i) loss of the RNA interference pathway components; ii) evidence of horizontal gene transfer events from bacteria; iii) the presence of genes unique to *Malassezia*; and iv) the expansion of secreted protein, lipase, and protease gene families that encode products predicted to breakdown lipids and proteins important for growth and host and microbial interactions.

The typical *Malassezia* genome is between ~7 and 9 Mb, which is about half the size of other basidiomycetous fungi, with the exception of *M. furfur* hybrid species whose genomes are twice the size of other *Malassezia* species. It is likely that the genomes of *Malassezia* species have reduced over time concomitantly with their evolution as a commensal organism and adaptation to the skin (Wu *et al*. 2015). There are other cases in which fungal genome reduction correlates with niche specialization, with the most remarkable examples being the obligate *Pneumocystis* species with genomes of ~7-8 Mb (Ma *et al*. 2016), and *Microsporidia* species with genomes as small as 2.9 Mb (Cuomo *et al*. 2012).

Aside from their commensal lifestyle, *Malassezia* fungi have been associated with several skin disorders, including pityriasis versicolor, dandruff, severe atopic dermatitis in humans, and otitis in dogs (Gaitanis *et al*. 2012; Wu *et al*. 2015). However, the exact role of *Malassezia* in these clinical conditions has been controversial, with recent studies even hypothesizing a protective role of *M. globosa* against *Staphylococcus aureus*, a bacterium that is associated with severe atopic dermatitis (Li *et al*. 2017; Ianiri *et al*. 2018). The lack of knowledge regarding *Malassezia* function within the skin mycobiome is due, in part, to the dearth of experimental systems for studying *Malassezia*-host interactions; current knowledge is based solely on in vitro experiments with isolated host cells (Watanabe *et al*. 2001; Ishibashi *et al*. 2006; Donnarumma *et al*. 2014; Glatz *et al*. 2015; Sparber and Leibundgut-Landmann 2017). Recently, 2 groundbreaking studies reported novel experimental murine models for studying *Malassezia* interactions with the skin and intestinal mucosa (Limon *et al*. 2019; Sparber *et al*. 2019). Sparber and colleagues demonstrated that the application of *M. sympodialis, M. pachydermatis*, and *M. furfur* on the dorsal ear skin of mice resulted in robust colonization of the epidermis and a rapid cytokine response dominated by IL-17 and related factors. This response was found to be critical for preventing fungal overgrowth on *Malassezia*-exposed skin and exacerbates inflammation under atopy-like conditions (Sparber *et al*. 2019). Another study by Limon and colleagues demonstrated the involvement of *Malassezia* in inflammatory bowel disease. The authors characterized the mycobiome associated with the intestinal mucosa of healthy individuals and patients with Crohn’s disease, and found that *M. restricta*, one of the most common inhabitants of human skin, was especially abundant in Crohn’s disease patients. Moreover, the presence of *M. restricta* was linked with a polymorphism in the gene for CARD9, a signaling adaptor critical for defense against fungi (Limon *et al*. 2019). The importance of these studies has been highlighted in 2 commentaries (Dawson 2019; Wrighton 2019).

Although these models represent an important advance in understanding the mechanisms of host responses to *Malassezia*, a lack of technologies for functional genetic studies has hampered the identification and characterization of the fungal components that promote inflammation and induce host responses. We were the first group to develop a transformation system based on transconjugation-mediated by *Agrobacterium tumefaciens* (AtMT, *A. tumefaciens*-mediated transformation) that is effective for both insertional and targeted mutagenesis and enabled the first genetic manipulation of *M. furfur* and *M. sympodialis* (Ianiri *et al*. 2016; Ianiri *et al*. 2017). Subsequently, *M. pachydermatis* has also been transformed (Celis *et al*. 2017).

CRISPR (clustered regularly interspaced short palindromic repeats)/Cas9 was originally discovered as a mechanism of adaptive bacterial immunity for defense against invading DNA elements (Jinek *et al*. 2012). The CRISPR/Cas9 system has been modified for use in other organisms, and at present, represents a revolutionary technology that has allowed gene editing in a number of cell types, including fungi (Shi *et al*. 2017; Adli 2018). The system consists of 2 elements: a specific endonuclease (Cas9) and a guide RNA (gRNA) that form a complex that catalyzes double-strand breaks (DSBs) at a specific DNA site flanking a protospacer adjacent motif (PAM) sequence of the host genome. After the DSB is generated, the DNA can be repaired either through non-homologous end joining (NHEJ) or through homology directed repair (HDR) when donor DNA is provided (Shi *et al*. 2017).

The present study is divided into 2 sections. In the first, we build upon the previously developed AtMT technology to perform the first T-DNA-mediated genetic screen in *M. furfur*. The aim was to generate a library of random insertional mutants, select for mutants with a phenotype of interest, and characterize insertion sites within the *M. furfur* genome to infer the function of genes involved in processes of physiological and clinical interest. In the second part of this study, we developed the first efficient, transient CRISPR/Cas9 mutagenesis system for *Malassezia*, and successfully generated targeted deletion mutants of two genes identified in the forward screen: *CDC55*, which encodes a subunit of protein phosphatase 2A (PP2A), and *PDR10*, which encodes an ABC transporter predicted to be involved in pleiotropic drug resistance. When validating the effectiveness of the T-DNA insertional mutagenesis for gene function studies, this novel CRISPR/Cas9 technology overcomes issues related to the reduced rate of homologous recombination observed in *M. furfur*, and we expect that it will facilitate molecular research on *Malassezia* fungi.

## Materials and Methods

### Strains and culture conditions

The haploid *M. furfur* strain CBS14141 (previously known as JPLK23) was used as the wild type (WT) strain for transformation experiments. This strain was maintained on modified Dixon’s media (mDixon) [mycological peptone (10 g/L), malt extract (36 g/L), glycerol (2 ml/L), tween 60 (10 ml/L), desiccated ox-bile (10 g/L) and agar (20g/L) for solid media]. Transformants were maintained on mDixon supplemented with the antifungal agents nourseothricin (NAT) or G418 (NEO).

### Forward genetics screen in *M. furfur*

Insertional mutagenesis was performed through AtMT using *Agrobacterium tumefaciens* strain EHA105 engineered with the binary vectors pAIM2 or pAIM6, which contain *NAT* and *NEO* resistance markers under the control of *M. sympodialis ACT1* promoter and terminator, respectively (Ianiri *et al*. 2016). Initially, transformations were performed using previously developed methods (Ianiri *et al*. 2016; Celis *et al*. 2017). Selected transformants were colony-purified on selective media and arrayed in 96 well plates containing 100 µL of mDixon + NAT or mDixon + NEO for in vitro assays and long-term storage.

For the primary screen, a 1.5 µL aliquot of cellular suspension of transformants was spotted on mDixon agar containing the following chemicals: Congo red (0.5%), sodium chloride (NaCl, 1M), sodium dodecyl sulfate (SDS, 0.3%), or fluconazole (FLC, 150 µg/ml) for cell wall and plasma membrane stress; NaNO_2_ (100 mM) for nitrosative stress; or CdSO_4_ (30 µM) for protein-folding defects and heavy metal stress. Transformants were also exposed to UV light (250 to 450 µJ × 100), elevated temperature (37°C), pH (pH 7.5), and nutrient-limiting conditions [yeast nitrogen base media (YNB)]; when used, arginine and tyrosine were added at 30 mg/L. Transformants selected in the primary screen as having a phenotype different than the WT were confirmed through a standard 1:10 serial dilution method by spotting 1.5 µL of cellular suspension on mDixon agar in the conditions that allowed their selection.

### Molecular characterization of the T-DNA insertional mutants of *M. furfur*

Insertional mutants with a phenotype of interest were single-colony purified and grown overnight in 25 mL of liquid mDixon for genomic DNA extraction using a CTAB extraction buffer (Pitkin *et al*. 1996). To identify the insertion sites of the T-DNA in the *M. furfur* genome, inverse PCR (iPCR) was performed according to previously published methods (Idnurm *et al*. 2004; Ianiri and Idnurm 2015). Briefly, approximately 2 µg of DNA were digested with the restriction enzymes PvuII, XhoI, SacII, ApaI, EcoRI (6-bp recognition site) or TaqI (4-bp recognition site), column purified, and eluted in 30 µL of elution buffer. Then, 8.5 µL of digested DNA were self-ligated with T4 DNA ligase (New England Biolabs) overnight at 4°C, and 1 µL was used as template for iPCR using primers ai76-ai77 for DNA digested with restriction enzymes that cut outside the T-DNA region, or ai076-M13F and ai077-M13R where restriction enzymes that cut inside the T-DNA were used (Idnurm *et al*. 2004). iPCR conditions were: initial denaturation at 94°C for 2 min, denaturation at 94°C for 30 sec, annealing at 55°C for 30 sec, and extension at 72°C for 2.5 min. PCR reactions were performed using ExTaq polymerase (Taqara Bio, Japan) according to manufacturer’s instructions. When ExTaq PCRs were unsuccessful, LaTaq polymerase (Taqara Bio, Japan) suitable for high G+C rich regions was used with an annealing temperature of 55°C and 60°C. Amplicons were either PCR- or gel-purified and subjected to Sanger sequencing. Sequences were subjected to BLASTn analysis against the *M. furfur* CBS 14141 genome assembly available on NCBI (reported as JPLK23) (Wu *et al*. 2015) and against an unpublished PacBio assembly. Gene boundaries and regulatory regions were determined using unpublished RNAseq data, which allowed us to define the accurate locations of T-DNA insertions. Retrieved *M. furfur* sequences were subjected first to BLASTx analysis against the latest genome assembly of *M. sympodialis* (Zhu *et al*. 2017) and subsequently on SGD (Saccharomyces Genome Database) to identify orthologs and infer gene function. Genes were named based on orthologous genes in *Saccharomyces cerevisiae*. Gene annotation was carried out manually based on BLAST searches and with the automated software Augustus (http://bioinf.uni-greifswald.de/augustus/submission.php) using RNAseq for untranslated regions (UTRs) and introns.

For Southern blot analysis, ~2 µg of genomic DNA were digested with SacII (no cut sites are within the *NAT* or *NEO* cassette, thus allowing us to determine the number of T-DNA insertions), resolved on a 0.8% agarose gel in 1x Tris-acetate EDTA (TAE) buffer, transferred to a Zeta-Probe membrane, and probed with *NAT* or *NEO* cassettes labeled with [^32^P]dCTP. *NAT* and *NEO* cassettes were amplified from plasmids pAIM2 and pAIM6, respectively, with universal M13F and M13R primers.

RNA extraction was performed using the standard TRIzol method (Rio *et al*. 2010). RNA was treated with the TURBO DNAse enzyme (Thermo Fisher Scientific) according to the manufacturer’s instructions, and quality was assessed using a NanoDrop spectrophotometer. Then, 3 µg of purified RNA were converted into cDNA via the Affinity Script QPCR cDNA synthesis kit (Agilent Technologies) according to manufacturer’s instructions. For each sample, cDNA synthesized without the RT/RNAse block enzyme mixture was used as a control for genomic DNA contamination. Approximately 500 pg of cDNA were used to measure the relative expression level of target genes through quantitative real-time PCR (RT-qPCR) using the Brilliant III ultra-fast SYBR green QPCR mix (Agilent Technologies) in an Applied Biosystems 7500 Real-Time PCR System. For each target, a “no-template control” was performed to analyze melting curves and to exclude primer artifacts. Technical triplicates and biological triplicates were performed for each sample. Gene expression levels were normalized using the endogenous reference gene *TUB2* and determined using the comparative ΔΔCt method.

### Generation of plasmid for CRIPSR/Cas9 targeted mutagenesis in *M. furfur*

Plasmids for targeted mutagenesis of *M. furfur CDC55* and *PDR10* through *A. tumefaciens*-mediated transformation were assembled in *S. cerevisiae* using the binary vector pGI3 as previously reported (Ianiri *et al*. 2016; Ianiri *et al*. 2017). The *NAT* cassette was amplified from plasmid pAIM1 using primers JOHE43277 and JOHE43278. The 5′ and 3′ flanking regions for homologous recombination were amplified from the genomic DNA of *M. furfur* CBS14141 using primer pairs JOHE45209-JOHE45210 and JOHE45211-JOHE45212 for *CDC55* and JOHE45201-JOHE45212 and JOHE45203-JOHE45204 for *PDR10*, respectively. The PCR products and the double-digested (KpnI and BamHI) binary vector pGI3 were transformed into S*. cerevisiae* using lithium acetate and PEG 3750 as previously reported (Ianiri *et al*. 2016). To assess correct recombination of the newly generated plasmids, single colonies of *S. cerevisiae* transformants were screened by PCR using primers specific for the *NAT* marker (JOHE43281–JOHE43282) in combination with primers homologous to outside of the region of the plasmid pGI3 involved in the recombination event (JOHE43279-JOHE43280). Positive clones of *S. cerevisiae* were grown ON in YPD and subjected to phenol-chloroform-isoamyl alcohol (25:24:1) plasmid extraction using a previously reported protocol (Hoffman 2001). The plasmid DNA obtained was then introduced into the *A. tumefaciens* EHA105 strain by electroporation, and the transformants were selected on LB + 50 µg/mL kanamycin. PCRs were performed using ExTaq and/or LATaq polymerase as described previously, with the only difference being an extension time of 1.5 min.

To generate the components of the CRISPR/Cas9 system in *Malassezia*, the histone H3 was identified in the *M. sympodialis* ATCC42132 genome assembly (Zhu *et al*. 2017) through BLASTp analysis using *S. cerevisiae* H3 as query. The 813-bp upstream and 257-bp downstream regions, including the *M. sympodialis* H3 promoter and terminator (indicated as p*H3*, and t*H3*), respectively, were amplified by PCR using JOHE46457-JOHE46458, and JOHE46461-JOHE46462, respectively. High Fidelity (HF) Phusion Taq polymerase (New England Biolabs) was used according to manufacturer’s instructions, with an annealing temperature of 55°C for 30 sec, and 1 min extension at 72°C. Primer JOHE46457 includes a chimeric region for recombination in pPZP-201BK and a multicloning site, and primers JOHE46458 and JOHE46461 include chimeric regions for recombination with primers JOHE46459 and JOHE46460, which were used to amplify *CAS9* open reading frame (ORF) from plasmid pXL1-Cas9 (Fan and Lin 2018). Primer JOHE46462 has SacII and SpeI restriction sites, and a region for recombination with the promoter of the 5SrRNA of *M. sympodialis* used to drive expression of the single guide RNA (gRNA). *CAS9* amplification did not work well with HF Phusion Taq, so we used ExTaq polymerase as described above, but with fewer cycles (20 cycles) and a 4-min extension. The *M. sympodialis* ATCC 42132 ribosomal cluster was identified in the latest genome assembly and annotation (Zhu *et al*. 2017) through BLASTn analysis using ITS sequences from *M. sympodialis* CBS 7222 available on GenBank (accession number NR_103583). A 674-bp region from the end of the rRNA-eukaryotic large subunit ribosomal RNA (position 612351 on chromosome 5), including the rRNA-5S ribosomal RNA gene (position 613025 on chromosome 5), was amplified by PCR using primers JOHE46463-JOHE46464. Primer JOHE46463 has a chimeric region complementary to primer JOHE46462. This PCR was performed using the touchdown protocol, with an initial denaturation of 94°C per 5 min, followed by 24 cycles of denaturation at 94°C for 30 sec, annealing at a gradient temperature of 62°C for 30 sec minus 1°C per cycle, and extension at 72°C for 1 min. This was followed by 16 cycles of denaturation at 94°C for 30 sec, annealing at 50°C for 30 sec, extension at 72°C for 1 min, with a final extension of 72°C for 5 min. The gRNA scaffold was amplified from plasmid pSDMA64 (Arras *et al*. 2016) using primers JOHE46465-JOHE46466. JOHE46466 includes SpeI and SacII restriction sites, 7 thymine residues (6T terminator), and a chimeric region for recombination in pPZP-201BK.

The specific target sequence for *CDC55* was identified using the program EuPaGDT (http://grna.ctegd.uga.edu/) available on FungiDB (https://fungidb.org/fungidb/). Specific target sequence for the gene *CDC55* was added by PCR with primers JOHE46468-JOHE46466 using the gRNA scaffold as template. Primer JOHE46468 has a chimeric region for recombination with both the 5SrRNA sequence and the gRNA scaffold, with an intervening target sequence specific for *CDC55*. These PCRs were performed using HF Phusion Taq as reported above. All components were gel purified, and equimolar amounts of the purified amplicons were used for overlap PCR to generate the Cas9 expression cassette (p*H3-CAS9*, t*H3*) and the complete gRNA (5S rRNA promoter fused with the gene-specific gRNA scaffold). PCRs were carried out using HF Phusion taq and the touch down protocol as above, with the only difference being extension times of 5 min and 1 min, respectively. The 2 resulting amplicons were cloned within the T-DNA of pPZP201BK digested with KpnI and BamHI through HiFi (New England Biolabs) assembly according to manufacturer’s instructions and recovered in *Escherichia coli* DH5α. *E. coli* clones were screened for recombinant plasmids by PCR using primers specific for the plasmid backbone (JOHE43279 and JOHE43280) in combination with JOHE46458 and JOHE46463, respectively. The plasmid sequence for CRISPR/Cas9 deletion of *CDC55* (named pGI40) was confirmed by Sanger sequencing.

To generate the CRISPR/Cas9 plasmid for targeted mutagenesis of *PDR10*, the binary vector pGI40 was digested with SpeI to remove the *CDC55*-specific gRNA, recovered from the gel and purified. A *PDR10*-specific target sequence designed using EuPaGDT was added to the gRNA scaffold by PCR using primers JOHE46466-JOHE46467, which have chimeric regions for recombination with both the 5SrRNA sequence and the gRNA scaffold, with the specific target sequence for *PDR10* in between.

This amplicon and the 5SrRNA previously generated were recombined through HiFi assembly within the T-DNA of the *Spe*I-digested pGI40, and the novel recombinant plasmids with the *PDR10*-specific gRNA (named pGI48) were identified by Sanger sequencing. This procedure is reported in Figure 4B. Recombinant plasmids were introduced in *A. tumefaciens* through electroporation.

AtMT was performed with modifications that increase transformation efficiency compared to our previous protocol used to generate insertional mutants. Briefly, *M. furfur* was grown for 2 days at 30°C and the culture was diluted to OD_600_ ~1. The engineered *A. tumefaciens* strains with the gene deletion cassettes and the CRISPR/Cas9 expression system were grown overnight, diluted to an OD_600_ ~0.1, and incubated for 4 to 6 h in shaking cultures (30°C) in liquid induction medium (IM) until OD_600_ reached a value of 0.6 to 0.8. These bacterial cellular suspensions were mixed in 1:1, 1:2, and 2:1 ratio, respectively, and they were added to *M. furfur* cellular suspension at 1:2 and 1:5 ratios, respectively. These cellular suspensions were centrifuged at 5200 *g* for 15 min, the supernatants were discarded, and ~500 µl to 1 mL of these fungal and bacterial mixes were spotted directly onto nylon membranes placed on mIM agar containing 200 µM acetosyringone. These were coincubated for 5 days at room temperature (plates maintained without Parafilm) prior to transferring the dual cultures to mDixon supplemented with NAT (100 µg/mL) to select for fungal transformants and cefotaxime (350 µg/mL) to inhibit *Agrobacterium* growth.

*M. furfur* transformants resistant to NAT were colony purified and subjected to phenotypic and molecular characterization. Putative mutants for the *CDC55* gene were exposed to UV light (250 to 300 µJ × 100) to identify those with impaired growth according to the results of the forward genetic screen. For molecular analysis, 23 representative NAT resistant transformants sensitive to UV light were subjected to phenol-chloroform-isoamyl alcohol (25:24:1) DNA extraction, and the correct replacement of the target loci was assessed by PCR. Diagnostic PCRs to identify homologous recombination events for the *CDC55* gene were carried out with primers JOHE45213 or JOHE45874 in combination with specific primers for the *NAT* gene (JOHE43281 and JOHE43282, respectively), and with primers JOHE45215-JOHE45216 specific for the internal region of *CDC55*. To evaluate the overall rate of homologous recombination (HR) of the CRISPR/Cas9 system, a larger number of *cdc55*Δ candidate mutants were tested for sensitivity to hydroxyurea, which was found to be the stressor with the strongest phenotype. Similarly, putative *pdr10*Δ mutants were exposed to FLC (150 µg/mL) for phenotypic characterization, and transformants displaying impaired growth were subjected to DNA extraction for molecular characterization. Diagnostic PCRs were carried out using primers JOHE45205 and JOHE45206 alone and in combination with specific primers for the *NAT* gene (JOHE43281 and JOHE43282, respectively), and with primers JOHE45207-JOHE45208 specific for the internal region of *PDR10*. PCR analyses consisted of 34 cycles of denaturation at 94°C for 30 sec, annealing at 55°C for 30 sec, extension at 72°C of 1 min/kb, with an initial denaturation at 94°C for 2 min and a final extension at 72°C for 5 min. PCR analyses were performed using ExTaq (Takara) according the manufacturer’s instructions. To detect homologous recombination events at the 3′ region of *CDC55* and to amplify the full length *PDR10* gene, LATaq polymerase (Takara) with Buffer I was used. PCR for *CAS9* was carried out with ExTaq and the touchdown protocol with primers JOHE46459-JOHE46461. PCR for the gRNA was carried out with JOHE46465-JOHE46466 using ExTaq as reported above. All the primers used are listed in Table S1.

Phenotypic analysis of the target mutants was performed on mDixon agar by spotting 1.5 µL of 1:10 dilutions of each cellular suspension in the following conditions: UV (250 to 450 µJ × 100), hydroxyurea (50 mM), benomyl (50 µM), FLC (150 µg/mL), amphotericin B (AmB, 50 µg/mL), 5-flucytosine (5FC, 1 mg/mL), caspofungin (Caspo, 100 µg/mL), cyclosporin A (CsA) at 100 µg/mL both alone and in combination with 10 mM of lithium chloride, tacrolimus (FK506) at 100 µg/mL both alone and in combination with 10 mM of lithium chloride, and dimethyl sulfoxide (DMSO, 40 µL) which was used to dissolve benomyl (10 mM).

### Genomic comparison and phylogeny of the ABC transporter of *M. furfur* and *M. sympodialis*

The predicted amino acid sequences of *S. cerevisiae* Pdr10, Pdr5, Pdr15, Pdr12, Snq2, Pdr18, Aus1, and Pdr11 were used as queries for tBLASTn and BLASTp searches against the genomes of *M. furfur* CBS14141 and *M. sympodialis* ATCC 42132 available on GenBank (Wu *et al*. 2015; Zhu *et al*. 2017). The *Malassezia* best hits were retrieved, and the encoded proteins for *M. furfur* were predicted using Augustus (http://bioinf.uni-greifswald.de/augustus/submission.php) based on RNAseq evidence for UTR regions and introns.

The web portal of ACT Artemis (https://www.webact.org/WebACT/home) was used for synteny analysis of a ~15000 bp region of *M. sympodialis* and *M. furfur* containing orthologues of the Pdr10 encoding genes. tBLASTx analysis with a E-value of 0.100000 was performed.

For phylogeny, the aforementioned predicted ABC transporter proteins were aligned using MUSCLE and the phylogenetic tree was generated with MEGA 7 (http://www.megasoftware.net/) (Kumar *et al*. 2016) using the maximum likelihood method (LG model, 5 discrete gamma categories) and 100 bootstrap replications.

## Results

### Molecular and phenotypic characterization of *M. furfur* insertional mutants

Insertional mutants of *M. furfur* were generated through AtMT using both *NAT* and *NEO* dominant drug resistance markers. A total of 767 insertional mutants were isolated and their growth was tested under several different stress conditions. A total of 19 mutants (~2.5%) with a phenotype different than the WT were selected for further characterization.

Inverse PCR (iPCR) was utilized to identify the genes inactivated by the T-DNA insertions. The sequenced amplicons were compared to the unannotated *M. furfur* CBS14141 genome assemblies (one reported in NCBI as *M. furfur* JPLK23, and another unpublished based on PacBio sequencing) coupled with RNAseq data, which facilitated the identification of the coding and regulatory sequences. This allowed an accurate determination of T-DNA insertion sites. Gene names were assigned according to the *Saccharomyces* Genome Database. Southern blot analysis was performed to determine the number of T-DNA insertions, revealing that 16 transformants harbored single T-DNA insertions, and 3 transformants had 2 T-DNA insertions (strains 4A10, 4B1, and 6B2) (Fig. 1). In parallel, iPCR allowed the characterization of 15 T-DNA insertions (Fig. 2). Six strains had T-DNA insertions within a predicted open reading frame (ORF), 2 strains had insertions within UTRs, and 3 strains had T-DNA insertions in intergenic regions with no RNAseq read coverage. Of the remaining 8 strains, 4 were suspected to have chromosomal rearrangements because the T-DNA borders were found in different locations in the *M. furfur* CBS 14141 genome, and the junctions between the T-DNA and the *M. furfur* genome could not be identified in another 4 strains. Table 1 summarizes the results of the forward genetics approach performed in this study, and properties of the T-DNA insertions are reported in Figure 2.

**Figure 1.**
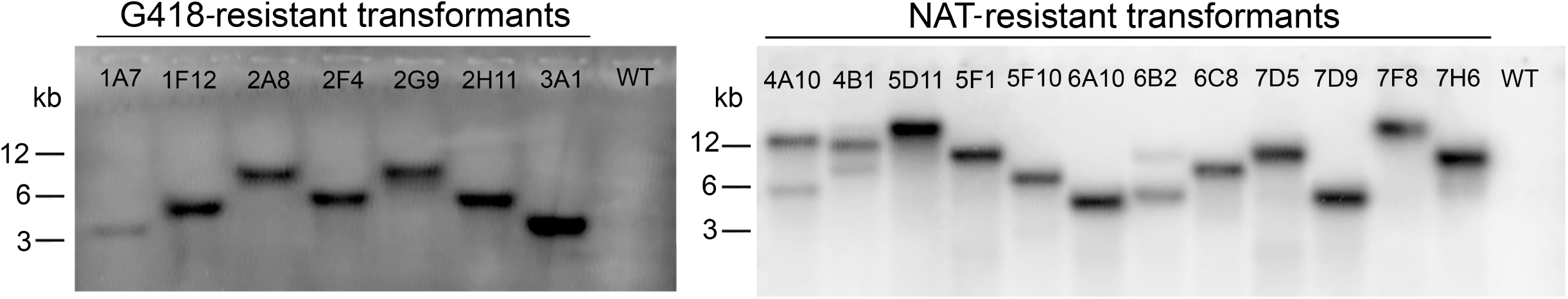
Southern blot analysis of *M. furfur* insertional mutants selected in the insertional genetic screen as having a phenotype different than the WT. Genomic DNA was digested with SacII, which does not cut within the *NAT* or *NEO* cassette, and hybridized with the ORF of the *NEO* (left) and *NAT* (right) genes. Each hybridization band corresponds to a single T-DNA insertion. The names of the transformants and of the *M. furfur* WT are indicated.

**Figure 2.**
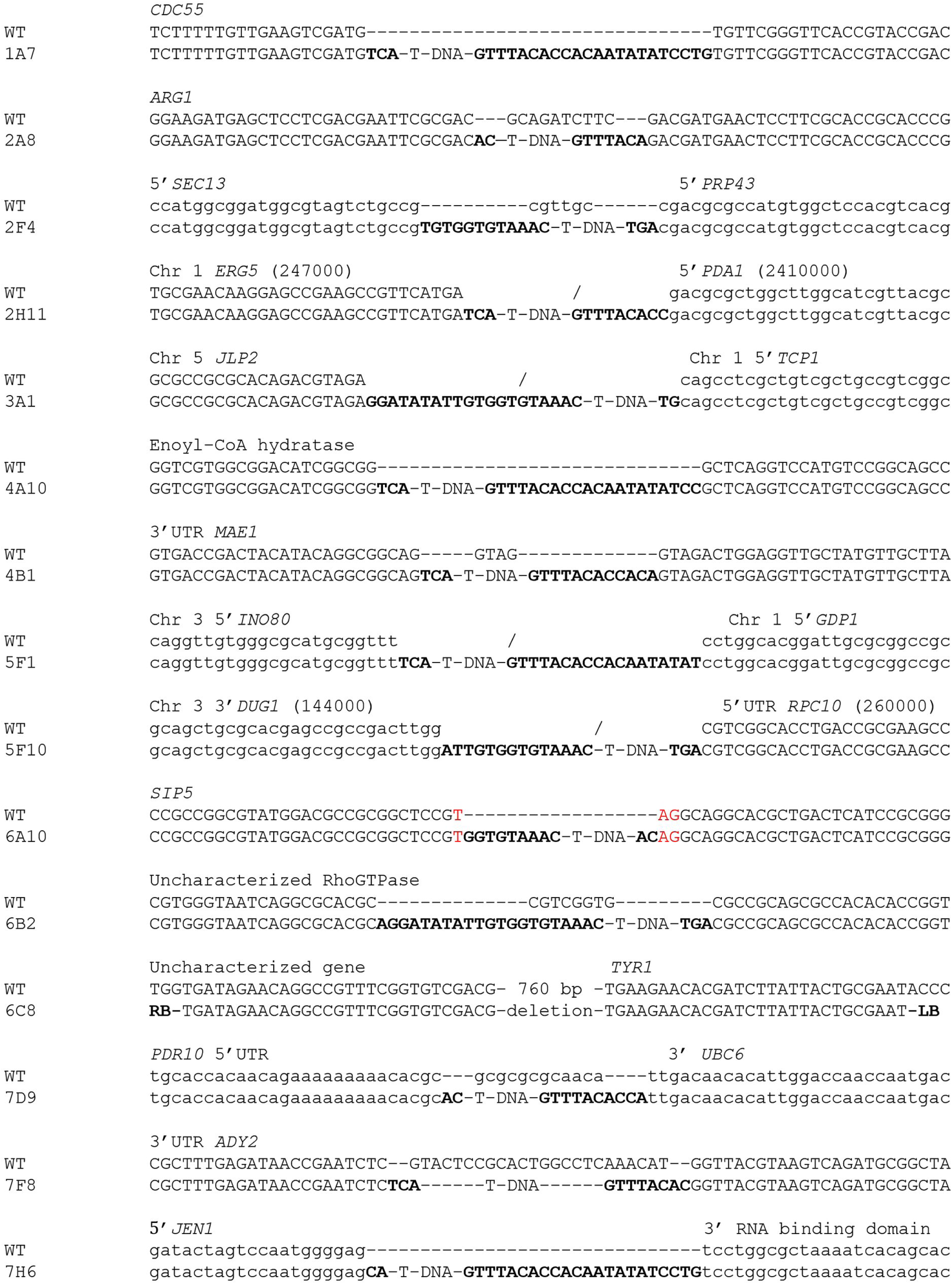
T-DNA insertion sites in 15 transformants of *M. furfur* as determined by iPCR. For each transformant, the mutated region and its corresponding region in the WT are shown. The region altered by the T-DNA is indicated above the sequence. When 2 regions are shown above the sequence, it indicates that the T-DNA insertion involved different locations of the *M. furfur* genome. The borders of the T-DNA are depicted in bold. Uppercase letters represent nucleotides corresponding to regions with RNAseq coverage (and hence representing either 5′ or 3′ UTRs as indicated or ORFs), while lowercase letters represent intergenic regions with no RNAseq coverage. The red nucleotides in strain 6A10 indicate a TAG stop codon. The symbol ‘-/-‘ indicates chromosomal rearrangement, with genomic locations shown in parentheses for the 2 transformants (2H11 and 5F10) having rearrangements involving the same chromosome.

**Table 1.**
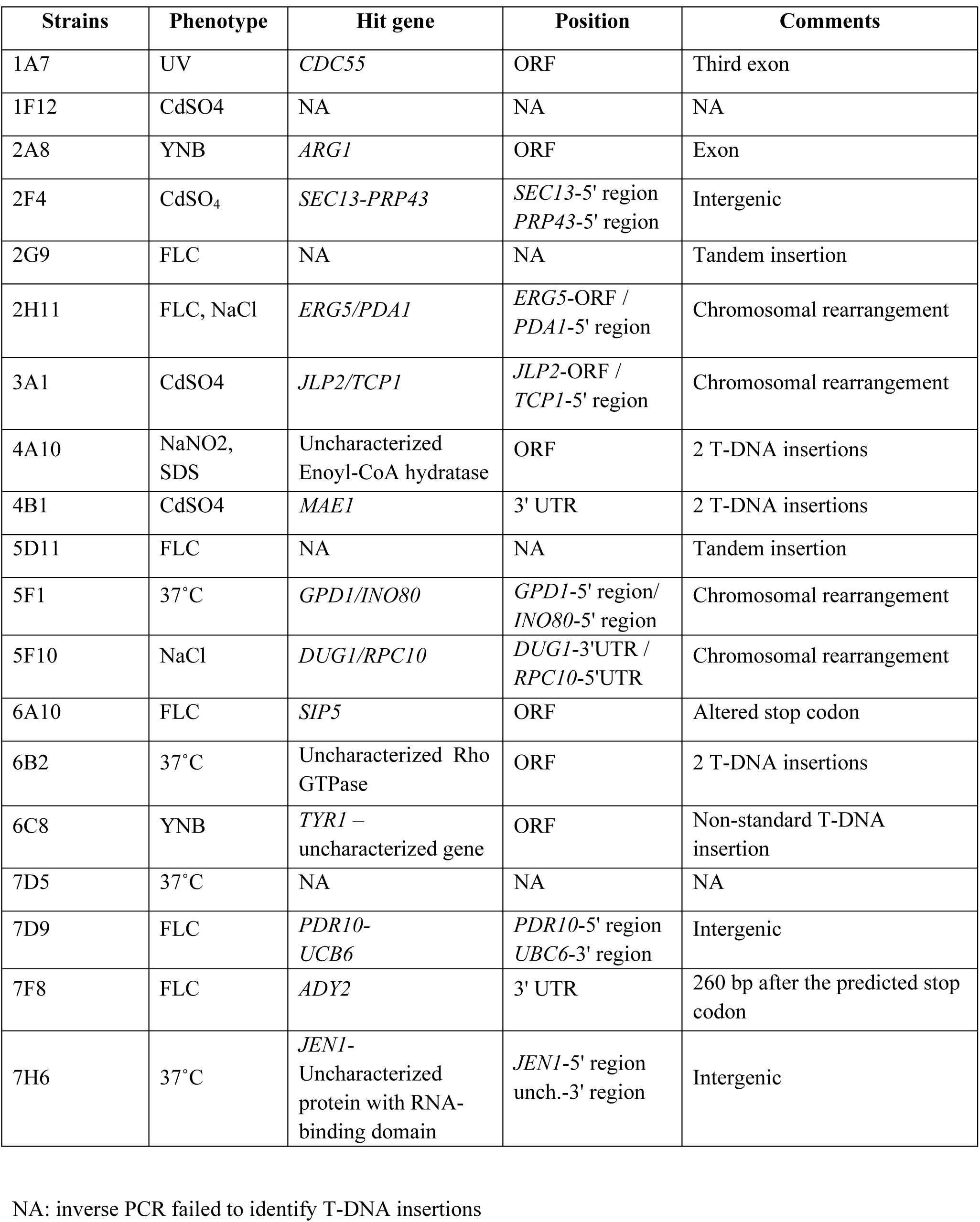
Insertional mutants for *M. furfur* isolated in the forward genetic screen.

Two mutants that displayed reduced growth on YNB were identified, and analysis of the genome sequence flanking the T-DNA revealed insertions in genes involved in amino acid biosynthesis (Fig. 3A). Strain 6C8 had a non-standard T-DNA insertion that generated a deletion of ~800 bp in the genome of *M. furfur*. Moreover, we were not able to identify the sequence of the left border (LB) from iPCR, and the first nucleotides obtained mapped within the ORF of *TYR1*, which encodes prephenate dehydrogenase, an enzyme involved in tyrosine biosynthesis (Mannhaupt *et al*. 1989). Conversely, the right border (RB) was found within the ORF of the adjacent gene encoding an uncharacterized protein with no conserved domains that shares similarity with several other *Malassezia* species and basidiomycetes. Addition of tyrosine did not restore the growth of strain 6C8 to the WT level (Fig. 3A), which instead was achieved in SD media supplemented with all amino acids, suggesting the hypothesis that Tyr1 is also involved in the biosynthesis of other amino acids. In strain 2A8, the T-DNA inserted within the ORF of *ARG1*, which encodes the enzyme arginosuccinate synthetase that catalyzes the formation of L-argininosuccinate from citrulline and L-aspartate in the arginine biosynthesis pathway (Jauniaux *et al*. 1978). Addition of L-arginine to YNB was sufficient to restore a WT phenotype, confirming that *M. furfur ARG1* is involved in arginine biosynthesis (Fig. 3 A).

**Figure 3.**
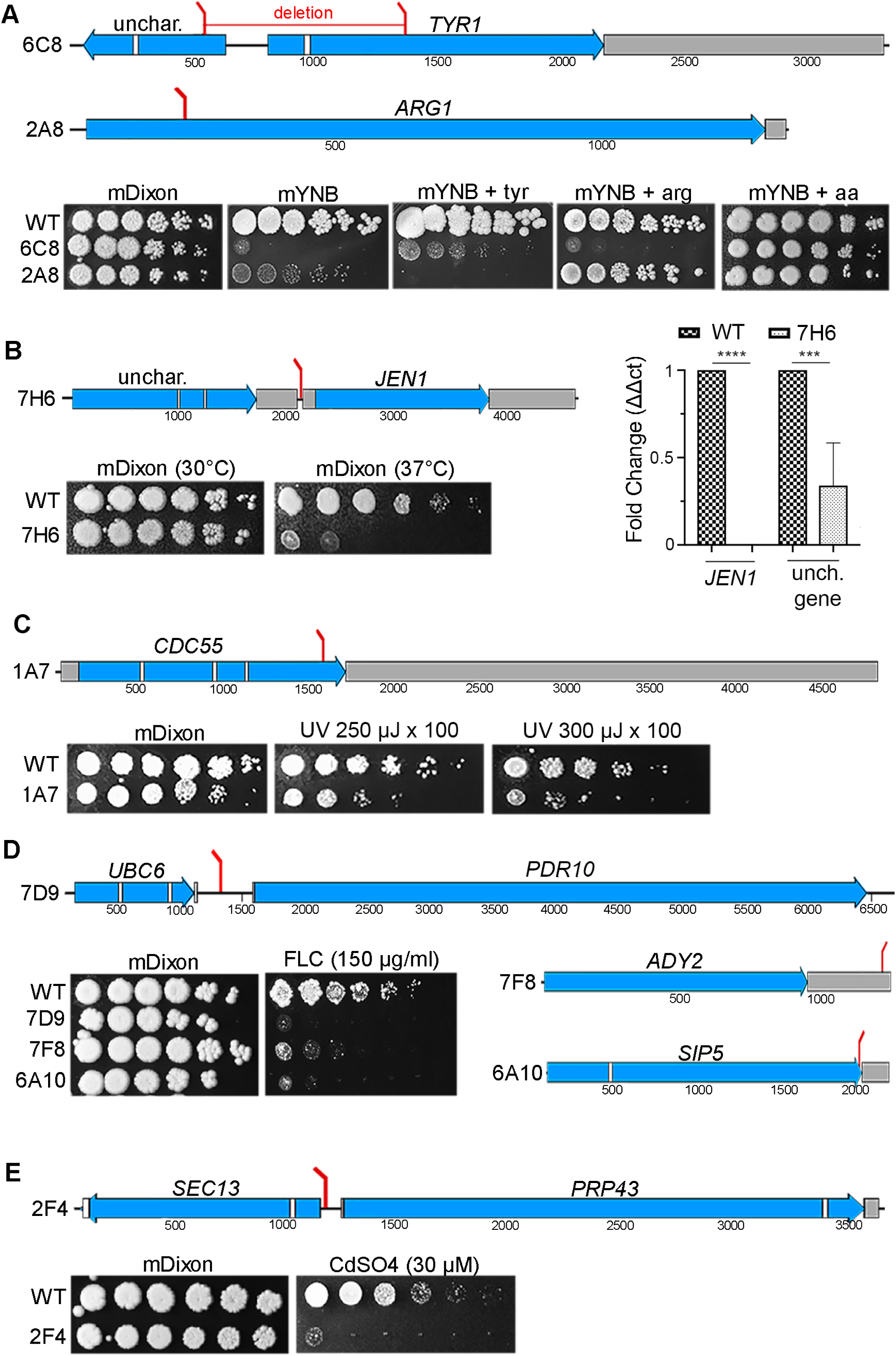
Position of T-DNA insertions in *M. furfur* mutants and their associated phenotypes. Each section shows mutants sensitive to the same stress or condition, such as reduced growth on minimal medium YNB (A), reduced growth at 37°C (B), and sensitivity to UV light (C), FLC (D), or cadmium sulfate (E). The positions of the T-DNA insertions are indicated by red bars, and in the same panel the phenotypes of the mutants are also shown. Exons are represented in blue, introns are in white, and UTRs are in gray. In the qPCR of panel 3B **** indicates p <0.0001 and *** indicates p <0.001 (p= 0.0008) for each pairwise comparison.

Four insertional mutants that showed decreased growth at elevated temperature (37°C) were identified. Of these, only strain 7H6 had a standard T-DNA insertion. In strain 7H6, the T-DNA integrated between 2 genes: downstream of an RNA-binding domain-containing protein and upstream of *JEN1* (Fig 3B). While the RNA-binding domain-containing protein is uncharacterized in *S. cerevisiae*, Jen1 is a plasma membrane monocarboxylate/proton symporter that transports pyruvate, acetate, lactate, and other substrates (Casal *et al*. 1999). To assess which gene was affected by the T-DNA insertion and therefore responsible for the phenotype of interest, an RT-qPCR analysis was performed. Expression levels were normalized to the *TUB2* gene of *M. furfur* WT grown at 30°C. Expression of the uncharacterized gene encoding the RNA-binding domain-containing protein in strain 7H6 was ~60% lower compared to the WT, whereas expression of *JEN1* was undetectable, indicating that either or both genes could be responsible for the temperature sensitive phenotype of strain 7H6 (Fig 3 B). The other transformants that displayed a temperature-sensitive phenotype included strain 5F1 (which showed a chromosomal rearrangement involving the 5′ regions of the gene *INO80* located on chromosome 1 and of the *GDP1* gene located on chromosome 3), strain 6B2 (which had 2 T-DNA insertions, one of which could be identified and was found within the ORF of a uncharacterized RhoGTPase), and strain 7D5 (whose T-DNA insertion could not be characterized by iPCR) (Table 1).

Strain 1 A7 showed increased sensitivity to UV light (250 and 350 µJ × 100) compared to WT *M. furfur* CBS 14141 (Fig. 3C). In strain 1A7, the T-DNA inserted into the third exon of the *CDC55* gene. In *S. cerevisiae, CDC55* encodes a regulatory subunit of protein phosphatase 2A. *CDC55* is involved in cell cycle control, and it is required for successful chromosome segregation and nuclear division (Healy *et al*. 1991; Bizzari and Marston 2011).

Six strains showed increased sensitivity to the antifungal fluconazole (FLC, 150 µg/ml) compared to the WT strain (Fig. 3D). Strain 6A10 had a T-DNA insertion in the predicted stop codon of the *S. cerevisiae* ortholog *SIP5*. The function of this protein is unknown, and it has no known domains. However, it has been reported to interact with both the Reg1/Glc7 phosphatase and the Snf1 kinase in response to glucose starvation (Sanz *et al*. 2000). In strain 7D9, the T-DNA was found in the intergenic region between the 5′ end of an ATP-binding cassette (ABC) multidrug transporter gene and the 3′ end of the *UBC6* gene. As shown in Figure 6E, the closest *S. cerevisiae* homolog is the ABC transporter *PDR10*, which is the designation that we adopted. ABC multidrug transporters are involved in pleiotropic drug responses that mediate resistance to xenobiotic compounds including mutagens, fungicides, steroids, and anticancer drugs (Sipos and Kuchler 2006). *UBC6* encodes a ubiquitin-conjugating enzyme involved in ER-associated protein degradation (Walter *et al*. 2001). As confirmed by targeted mutagenesis (discusses below), the FLC-sensitive phenotype of strain 7D9 is due to T-DNA insertion in the promoter region of *PDR10*.

In strain 7F8 the T-DNA inserted within the 3’UTR of *ADY2* (Fig. 3D). *ADY2* encodes an ammonium and acetate transmembrane transporter involved in nitrogen utilization (Rabitsch *et al*. 2001; Paiva *et al*. 2004). In strain 2H11 T-DNA integration generated a rearrangement involving the *ERG5* gene and the region close to the 5′ end of the *PDA1* gene. In addition to increased sensitivity to FLC, this strain showed increased sensitivity to sodium chloride compared to WT. Erg5 is a cytochrome P450 enzyme that is a C-22 sterol desaturase involved in ergosterol biosynthesis (Lees *et al*. 1995). Although it is known that FLC targets membrane ergosterol, and *ERG5* deletion in *S. cerevisiae* leads to increased FLC sensitivity (Kapitzky *et al*. 2010), it cannot be excluded that the FLC and NaCl sensitivity of strain 2H11 is due both to *ERG5* mutation and the intrachromosomal rearrangement itself. In strains 2G9 and 5D11, the T-DNA likely integrated in tandem repeats because iPCR amplicons consisted of both the left and right borders of the T-DNA fused together, and this prevented retrieval of the junctions between the T-DNA and the genome.

Three strains showed sensitivity to cadmium sulfate (CdSO_4_, 30 µM), and only one (strain 2F4) showed a standard T-DNA insertion in the 5′ regions of both *SEC13* and *PRP43* (Fig. 3E). *SEC13* in *S. cerevisiae* encodes an essential protein that is a structural component of the COPII (coat protein complex II), of the nuclear pore outer ring, and of the Seh-1 associated complex. It is involved in COPII-coated vesicle budding from the ER to the Golgi, nuclear pore distribution, and the ubiquitin-dependent ERAD (ER-associated ubiquitin-dependent protein breakdown) pathway, which is involved in protein degradation by cytoplasmic proteasomes (Menon *et al*. 2005; Dokudovskaya *et al*. 2011; ČopiČ *et al*. 2012). *S. cerevisiae PRP43* encodes an RNA helicase protein that is also essential for viability and contributes to the biogenesis of ribosomal RNA, and it is also involved in spliceosomal complex disassembly (Arenas and Abelson 1997; Giaever *et al*. 2002). qPCR did not show clear downregulation of either gene (data not shown), and whether either or both genes are responsible for the cadmium sulfate-sensitive growth defect remains to be established. Because the T-DNA inserted in the 5′ region of *SEC13* and *PRP43*, whose orthologs are essential in *S. cerevisiae*, we speculate that the functions of both genes are affected or that the phenotype observed is unlinked to the T-DNA insertion. In strain 3A1, a rearrangement involving the *JLP2* and *TCP1* genes was found, and for strain 4B1, Southern blot indicated 2 T-DNA insertions, one of which was identified and found within the 3’UTR of the *MAE1* gene, which encodes a mitochondrial malic enzyme that is important for sugar metabolism and acts as a precursor for many amino acids (Boles *et al*. 1998). For strain 1F12, iPCR using different restriction enzymes was unsuccessful. We also identified a strain (4A10) that was sensitive to sodium nitrite (NaNO_2_) and SDS. According to Southern blot analysis, strain 4A10 has 2 T-DNA insertions, one of which could be identified and was found in an uncharacterized enoyl-CoA hydratase gene. Another strain (5F10) was sensitive to NaCl and iPCR revealed the presence of a chromosomal rearrangement involving the 3′ region of the *DUG1* gene and the 5′ UTR of the *RPC10* gene.

### Development of a CRISPR/Cas9 gene deletion system to generate *cdc55*Δ *M. furfur* mutants

To validate the results of the insertional mutagenesis screen, the insertional mutants 1A7 and 7D9 and their mutated genes were chosen for further analysis as a proof of principle. First, we focused on the UV-sensitive strain 1A7 with a T-DNA insertion in the *CDC55* gene. We were intrigued by this strain because *CDC55* mutation is not known to be responsible for UV sensitivity in other fungi. The aim was to generate an *M. furfur cdc55*Δ targeted mutant, determine if the UV phenotype of the original insertional mutant is attributable to *CDC55* mutation, and investigate any further functions of the gene in *M. furfur*.

For targeted mutagenesis of *CDC55*, molecular biology techniques were performed following our previously published methods (Ianiri *et al*. 2016). Regions of 1500 and 1000 bp flanking the 5′ and 3′ ends of the *CDC55* target gene, respectively, were amplified from *M. furfur* genomic DNA and fused with the *NAT* marker within the T-DNA borders of plasmid pGI3. The recombinant plasmid (pGI41) bearing the *cdc55*Δ*::NAT* allele was identified in *S. cerevisiae* by colony PCR and *A. tumefaciens* EHA105 transformed by electroporation. Several rounds of *Agrobacterium*-transconjugation were performed, and NAT-resistant transformants of *M. furfur* were single colony-purified and subjected to diagnostic PCR to confirm *CDC55* targeted mutagenesis. None of the transformants tested (0 out of more than 100) showed full replacement of the gene *CDC55*.

Next we developed a CRISPR/Cas9 system for *M. furfur* to increase homologous recombination efficiency. Because the plasmid for targeted gene replacement of *CDC55* was already available, we generated an additional plasmid to make a DNA DBS in *CDC55*, and then used the available *cdc55*Δ*::NAT* allele as HDR template to repair the break. For expression of Cas9, the ORF of the *CAS9* endonuclease was cloned under the control of the strong promoter and terminator of the histone *H3* gene of *M. sympodialis* ATCC42132. To drive expression of gRNA specific for the target *CDC55* gene, the promoter of the 5S rRNA was chosen. Because the ribosomal cluster is well annotated in the newly released genome of *M. sympodialis* (Zhu *et al*. 2017) and we have evidence that *M. sympodialis* promoters and terminators are functional in *M. furfur* (Ianiri *et al*. 2016), a 689 bp region including the 5S rRNA and its upstream region was amplified from *M. sympodialis* ATCC42132. The forward primer for the p5S rRNA contained restriction sites for SacII and SpeI to facilitate genetic manipulations. The scaffold gDNA also was obtained by PCR, and 6 thymine residues (6-T) were included as terminator. A 20-nt oligonucleotide target of gRNA was designed to match a region of the *CDC55* gene adjacent to a PAM site, and it included the 5′ and 3′ regions that overlapped with the 5S rRNA promoter and the gRNA scaffold, respectively. This target oligonucleotide was added to the gRNA scaffold through PCR as reported in Figure 4B. The 5 PCR fragments (p*H3*; *CAS9*; t*H3*; p5S rRNA; gRNA) were used as template for overlap PCRs, and two final amplicons (p*H3*-*CAS9*-t*H3* and p5SrRNA-gRNA) were cloned in pPZP201BK to generate plasmid pGI40 (Table 2) (Figures 4A and 4B).

**Table 2.** Plasmid s used in the present study.

| Name | Background | Relevant features | Purpose | References |
| --- | --- | --- | --- | --- |
| pAIM2 | pPZP-201BK | p*ACT1-NAT-tACT1* | Random insertional mutagenesis | (IANIRI *et al.* 2016) |
| pAIM6 | pPZP-201BK | p*ACT1-NEO-tACT1* | Random insertional mutagenesis | (IANIRI *et al.* 2016) |
| pPZP-201BK | NA | KAN-R | Binary vector that replicates in *E. coli* and *A. tumefaciens* | (COVERT *et al.* 2001) |
| pGI3 | pPZP-201BK | Sc*URA3* + 2μ; KAN-R | Binary vector that replicates in *E. coli*, *A. tumefaciens* and *S. cerevisiae* | (IANIRI *et al.* 2017) |
| pGI40 | pPZP-201BK | p*H3-CAS9-tH3* + p5S rRNA-sgRNA *CDC55* | Cas9 endonuclease and *CDC55* target sgRNA | This study |
| pGI41 | pGI3 | *cdc55::NAT* | HDR template to generate *cdc55Δ* | This study |
| pGI42 | pGI3 | *pdr10::NAT* | HDR template to generate *pdr10Δ* | This study |
| pGI48 | pGI40 | p*H3-CAS9-tH3* + p5S rRNA-sgRNA *PDR10* | Cas9 endonuclease and *PDR10_1* target sgRNA | This study |

- Covert, S. F., P. Kapoor, M.-H. Lee, A. Briley and C. J. Nairn, 2001 *Agrobacterium tumefaciens*-mediated transformation of *Fusarium circinatum*. *Mycol. Res.* 105: 259-264. - Ianiri, G., A. F. Averette, J. M. Kingsbury, J. Heitman and A. Idnurm, 2016 Gene function analysis in the ubiquitous human commensal and pathogen *Malassezia* genus. *mBio* 7: e01853-01816. - Ianiri, G., K. J. Boyce and A. Idnurm, 2017 Isolation of conditional mutations in genes essential for viability of *Cryptococcus neoformans*. *Curr Genet* 63: 519-530.

**Figure 4.**
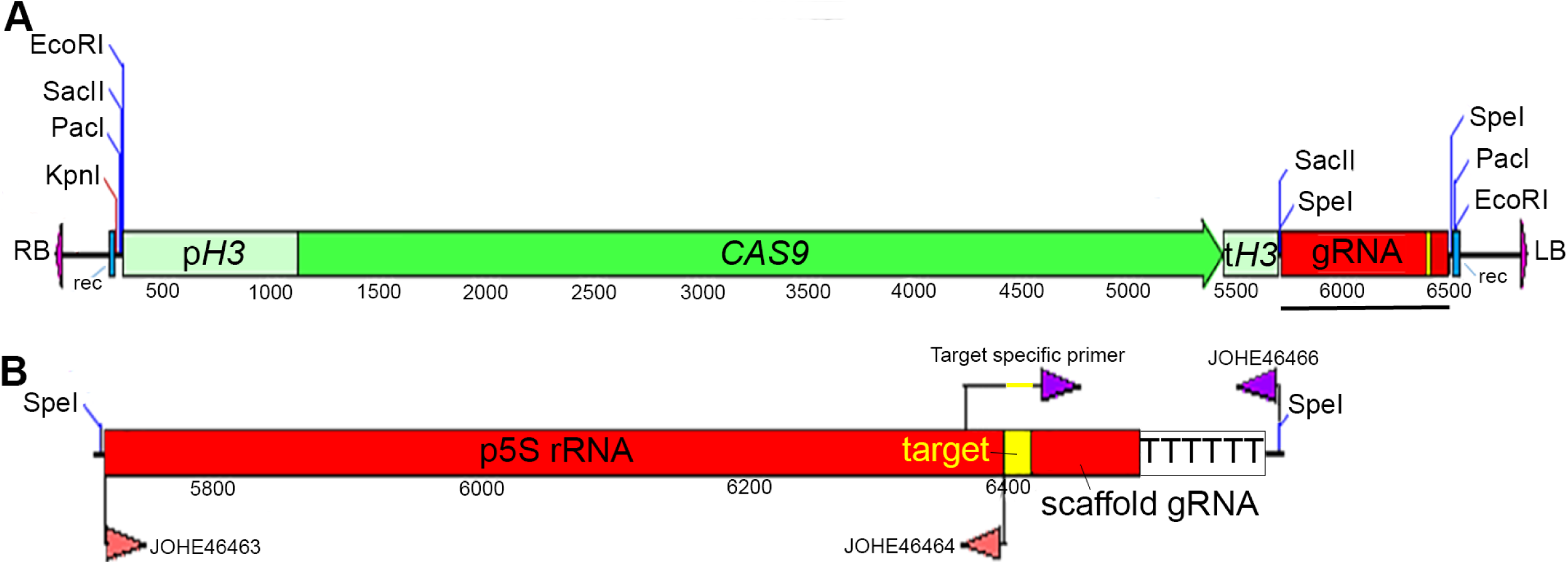
Development of a CRISPR/Cas9 system in *M. furfur*. (A) Complete T-DNA necessary for *CAS9* expression and gRNA excision in *M. furfur*. The promoter and terminator of the histone *H3* gene of *M. sympodialis* ATCC42132 (p*H3* and t*H3*, respectively) and the *CAS9* gene are shown in green. The gRNA is shown in red, and the gene-specific target region is shown in yellow. Sites for recombination (rec) are shown in blue, and the right and left borders (RB and LB, respectively) of the T-DNA are shown in purple. Restriction sites were added to facilitate further molecular manipulation. The black bar indicates the full-length gRNA that is shown in greater resolution in B. (B) The gRNA includes the 5S rRNA promoter region (p5S rRNA) obtained with primers JOHE46463 and JOHE46464 through touchdown PCR with *M. sympodialis* ATCC42132 genomic DNA. The gene-specific gRNA was obtained by PCR using a target-specific primer, which overlaps with the p5S rRNA and the gRNA scaffold and includes a 20 nt target sequence in between them (represented in yellow). This was used in combination with JOHE46466, which includes the 6T terminator (and also a region for recombination in pPZP201BK, which is not shown). A SpeI restriction site was added to facilitate further use of the CRISPR/Cas9 system in *Malassezia*. To perform targeted mutagenesis of another gene, we recommend using SpeI digestion of the plasmid in A (pGI40) and cloning the 2 PCR products (p5S rRNA and gene-specific gRNA) through Gibson or HiFi assembly. We used this strategy to generate binary vector pGI48 for CRISPR/Cas9-mediated mutagenesis of the *PDR10* gene.

AtMT of *M. furfur* CBS14141 was conducted to test the developed CRISPR/Cas9 system to generate targeted gene replacement of the *CDC55* gene. Since our previous reports of AtMT of *Malassezia* (Ianiri *et al*. 2016; Ianiri *et al*. 2017), we have optimized the protocol to achieve a higher transformation efficiency. The main change included the use of a 2:1 to 5:1 *Malassezia*:*A. tumefaciens* mixture that was concentrated through centrifugation before the coincubation step on modified induction media [mIM, (Ianiri *et al*. 2016)]. The detailed procedure is reported in the Materials and Methods. For co-transformation of *M. furfur* using *A. tumefaciens* strains bearing the binary vectors pGI40 (CRISPR/Cas9 expression system) and pGI41 (HDR *cdc55*Δ*::NAT* template), induced bacterial strains were mixed in ratios of 1:1, 1:2, and 2:1, then added to ratios of 1:2 and 1:5 with *M. furfur* cells (Fig. 5A). The co-cultures were centrifuged to eliminate the supernatants, and the pellet containing the mix of the 3 components was spotted on nylon membranes placed on mIM agar. The plates were incubated at room temperature for 5 days without parafilm. The coincubation cultures were recovered and plated on mDixon containing NAT and CEF. A representative subset of 23 *M. furfur* transformants was single colony-purified and subjected to molecular characterization through PCR. Genotyping was performed using 1) primers designed beyond the regions of DNA used in the generation of the deletion allele in combination with specific *NAT* primers; 2) primers internal to the gene *CDC55*; 3) primers specific for the *CAS9* genes, and primers specific for the gRNA (Figure 5B). Specific amplicons of ~1.6 kb and ~1.4 kb for the 5′ (left) and 3′ (right) T-DNA-genomic DNA junctions, respectively, were obtained for all of the 23-randomly selected transformants. Accordingly, the internal region of *CDC55* was amplified only from the WT strain. No amplicons for *CAS9* or the gRNA were obtained. These results indicate that all transformants tested had full replacement of the *CDC55* gene and absence of *CAS9* and gRNA integration in the genome (Fig. 5C). Furthermore, 64 additional random *cdc55*Δ mutant candidates were tested for sensitivity to hydroxyurea, which we found to be the most effective stressor for the *cdc55*Δ phenotype, and found that 62 displayed impaired growth compared to WT (Fig. S1). Therefore, molecular and phenotypic analyses revealed that out of 87 transformants analyzed, 85 were *cdc55*Δ mutants, resulting in a rate of homologous recombination of 97.7%.

**Figure 5:**
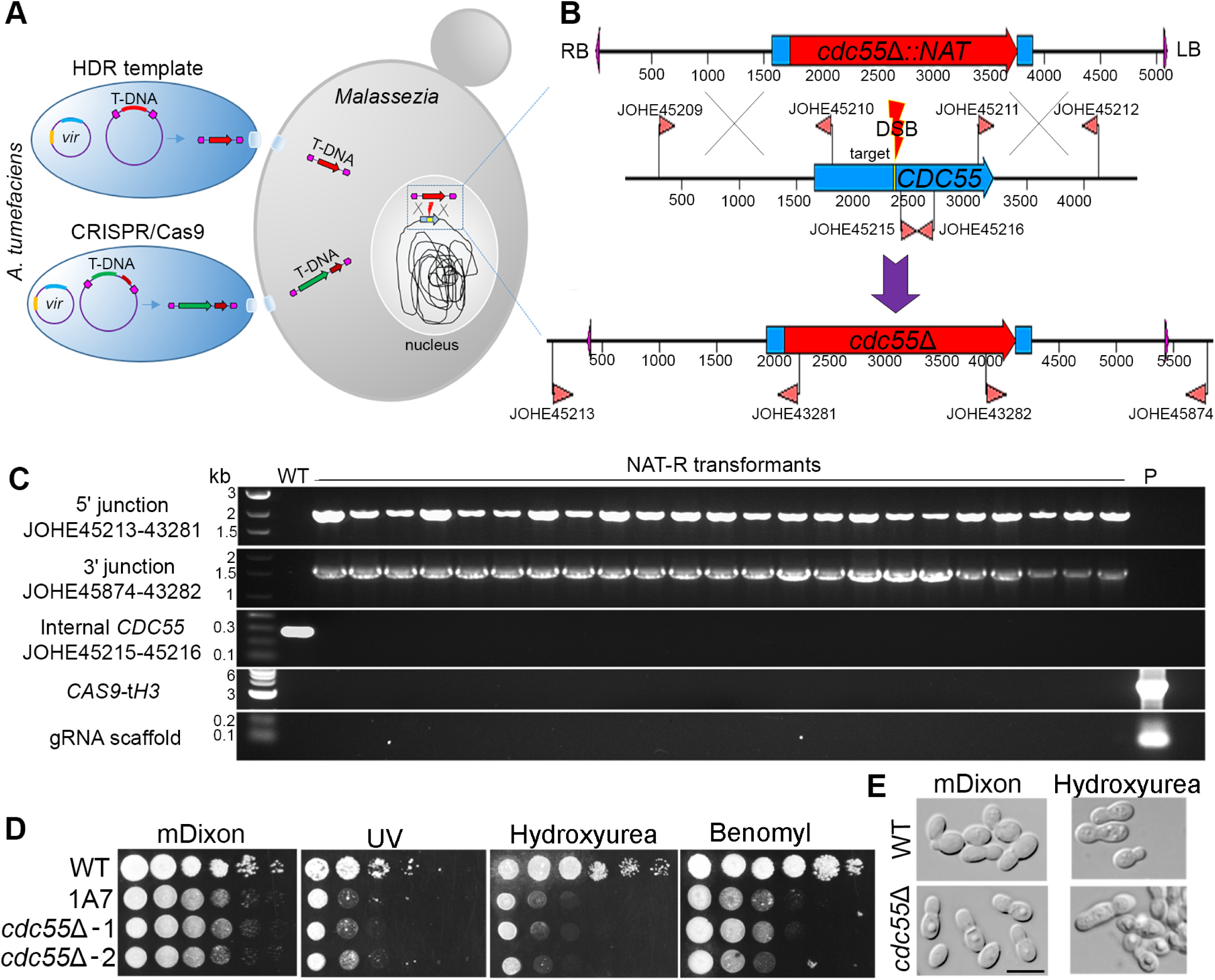
CRISPR/Cas9-mediated target mutagenesis of *M. furfur CDC55*. (A) Co-transformation of *M. furfur* mediated by 2 *A. tumefaciens* strains, one that delivers a T-DNA including the HDR template (ie *cdc55*Δ*::NAT* deletion construct in red) and another that includes the *CAS9* cassette (in green) and the gene-specific gRNA (dark red). Also depicted are the *vir* plasmids present in *A. tumefaciens* cells that are necessary for T-DNA excision and transfer to the *M. furfur* nucleus where homologous recombination occurs. (B) Magnification of the homologous recombination event that occurs in the *M. furfur* nucleus. The top construct represents the T-DNA of the plasmid pGI41 bearing the *cdc55*Δ*::NAT* HDR template. The middle panel represents the native *M. furfur CDC55* locus, the primers used to amplify the 5′ and 3′ regions for HR, the internal primer for the *CDC55* gene, and the 20-nt target sequence (yellow). The gRNA guides Cas9 to the target site to generate a DBS that is repaired using the deletion allele as template, resulting in the targeted replacement of *CDC55* with a *NAT* dominant marker (lower panel). Primers outside the region in which homologous recombination events occur are used in combination with primers for the *NAT* marker to identify *cdc55*Δ mutants. (C) Diagnostic PCR analyses of *M. furfur* WT and NAT-resistant transformants for the identification of *cdc55*Δ mutants. Each panel used the indicated combination of primers, whose position can be found in panel B. PCR primers for *CAS9* and gRNA are reported in Table S1. (D) Phenotypic analysis of *M. furfur* WT, insertional mutant 1A7, and two independent *cdc55*Δ mutants on mDixon (control), UV (300 µJ × 100), hydroxyurea (50 mM) and benomyl (50 µM); 1.5 µL of tenfold serial dilution were spotted on the agar plates, incubated at 30°C for 3 to 7 days, and then photographed. (E) Microscopic analysis of cell morphology of WT and a representative *cdc55*Δ mutant after growth on mDixon and mDixon supplemented with hydroxyurea (50 mM); the black bar indicates 5 µm.

In *S. cerevisiae, CDC55* positively regulates mitotic entry at the G2/M phase transition and negatively regulates mitotic exit, and it regulates the mitotic spindle assembly and the morphogenesis checkpoint (Wang and Burke 1997; Bizzari and Marston 2011). Null *cdc55*Δ mutants display abnormally elongated buds; decreased growth rate; and increased sensitivity to gamma rays and hydroxyurea (DNA-damaging agents that interfere with DNA replication), to benomyl and nocodazole (which interfere with microtubule polymerization), and cold-induced stress. Phenotypic characterization of 2 representatives independent *cdc55*Δ *M. furfur* mutants confirmed that they were sensitive to UV light (Fig. 5D), corroborating the phenotype of the insertional mutant 1A7. Moreover, the *M. furfur cdc55*Δ mutant had a slower growth rate compared to the WT strain, and increased sensitivity to hydroxyurea and benomyl (Fig. 5D). Due to the inability of *M. furfur* WT to grow at low temperature, cold sensitivity could not be determined for *M. furfur cdc55*Δ. *M. furfur cdc55*Δ mutants were subjected to microscopy analysis both under normal and stress conditions, and when exposed to hydroxyurea they displayed cells with abnormal morphology and elongated buds, similar to *S. cerevisiae cdc55*Δ mutants (Fig. 5E).

### Generation of a *pdr10*Δ *M. furfur* mutant with CRISPR/Cas9

The other insertional mutant of interest was strain 7D9, which has a T-DNA insertion between the *PDR10* and *UBC6* genes and exhibited FLC sensitivity. It was hypothesized that the phenotype of strain 7D9 was due to the T-DNA interfering with the function of *PDR10*, which is well known to mediate antifungal drug response and therefore was chosen for targeted mutagenesis. In *M. furfur*, *PDR10* is a large, 4470-bp gene. The *pdr10*Δ*::NAT* gene disruption cassette was generated as previously described for *CDC55*, and the vector was named pGI42. For the gRNA, a primer with a specific *PDR10* target between regions that overlap with the 5S rRNA promoter and the gRNA scaffold was added to the gRNA scaffold by PCR. The resulting amplicon was then cloned together with the p5S rRNA in the T-DNA of pGI40 digested with SpeI as reported in Figure 4B; this vector was named pGI48.

Co-transformation of *M. furfur* CBS14141 was performed using *A. tumefaciens* strains bearing plasmids pGI42 and pGI48 as reported in Figure 5A. 60 *M. furfur* NAT-R transformants were single colony purified, and streaked onto mDixon + FLC. Five (8.3%) transformants that displayed FLC sensitivity plus a randomly-selected FLC-resistant control strain were subjected to molecular characterization (Fig. 6B). PCR analysis using external screening primers designed beyond the region of DNA utilized to generate the *pdr10*Δ*::NAT* deletion allele produced 2 amplicons: a 6183-bp amplicon for the WT and the FLC resistant strain, and a 3933-bp amplicon for the 5 transformants that displayed FLC sensitivity. For these 5 transformants, PCR carried out using the external primers with specific *NAT* primers generated amplicons of ~1.1 kb and ~1.3 kb on the 5′ and 3′ regions, respectively. PCR using primers internal to the *PDR10* gene generated an amplicon of 486 bp only in the WT and the randomly selected NAT-R strain. These PCR results indicate full replacement of the *PDR10* gene in the 5 transformants that displayed FLC sensitivity.

**Figure 6.**
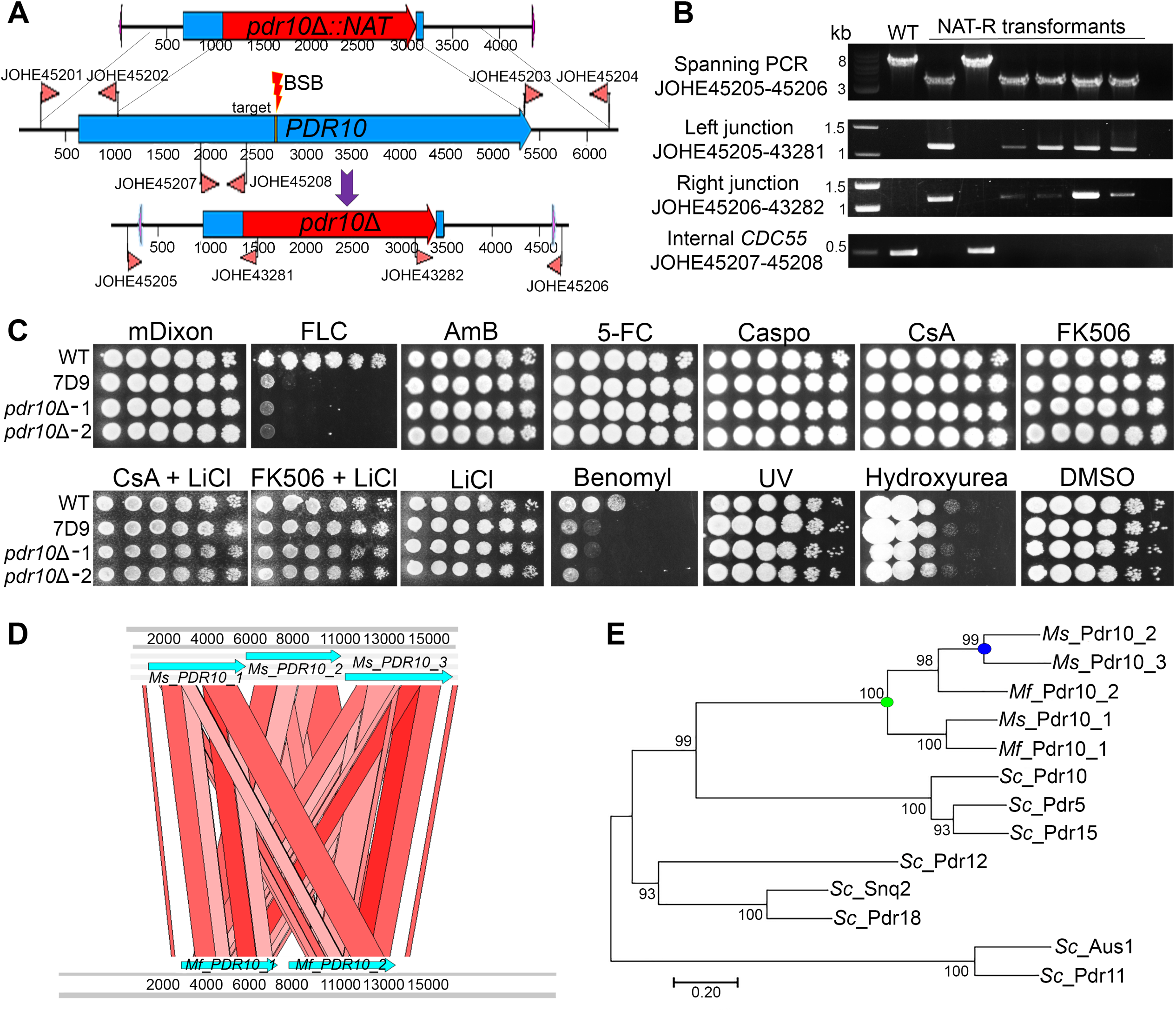
Targeted CRISPR/Cas9-mediated gene replacement of the *M. furfur PDR10* gene. (A) The T-DNA including the *pdr10*Δ*::NAT* HDR template is shown in the top panel; The *PDR10* gene, the primers used to amplify the 5′ and 3′ regions for homologous recombination, the internal primer for the *PDR10* gene, and the 20-nt target sequence (yellow) for the DBS are shown in the middle panel. The bottom panel shows the *pdr10*Δ mutant allele and the primers used for PCR. (B) Diagnostic PCR analyses of *M. furfur* WT and 5 FLC sensitive and 1 FLC-resistant transformants for the identification of *pdr10*Δ mutants. Each panel used a combination of primers that are represented in panel A. (C) 1.5 µl of cellular suspension of the WT strain CBS 14141, insertional mutant 7D9, and 2 independent *pdr10*Δ mutants were spotted on mDixon (control), FLC (150 µg/mL), amphotericin B (AmB, 50 µg/mL), 5-fluorocytosine (5-FC, 1 mg/mL), caspofungin (Caspo, 100 µg/mL), cyclosporine A alone or with 10 mM of LiCl (CsA, 100 µg/mL), FK506 (100 µg/mL) alone or with 10 mM of LiCl, benomyl (50 µM), UV (300 µJ × 100), hydroxyurea (50 mM), or dimethyl sulfoxide [DMSO, 800 µL/L (solvent used to resuspend benomyl)]. (D) ACT Artemis synteny comparison of a 15-kb region including the 3 copies of the *PDR10* gene of *M. sympodialis* (*Ms*) and the 2 copies of the *PDR10* gene of *M. furfur* (*Mf*). (E) The predicted proteins of the *S. cerevisiae* (*Sc*) ABC transporters Pdr10, Pdr5, Pdr15, Pdr12, Snq2, Pdr18, Aus1, and Pdr11; *M. sympodialis* (*Ms*) Pdr10_1, Pdr10_2, and Pdr10_3; and *M. furfur* (*Mf*) Pdr10_1 and Pdr10_2 were used to generate a maximum likelihood phylogenetic tree with the LG + G method (100 bootstrap replications).

Mutants *pdr10*Δ showed hypersensitivity to FLC, indicating that the phenotype of strain 7D9 was due to the T-DNA interfering with the function of *PDR10*. Moreover, because ABC transporters are known to be involved in pleiotropic drug resistance and cellular detoxification, the phenotypic response of *pdr10*Δ mutants was tested against other antifungal drugs of clinical relevance. Surprisingly, *M. furfur pdr10*Δ mutants showed only sensitivity to FLC and grew at the WT level on amphotericin B, 5-flucytosine, caspofungin, tacrolimus (FK506), and cyclosporine A (CsA) both alone and in combination with the plasma membrane stressor lithium chloride (Fig. 6C and data not shown), which we previously showed enhances antifungal activity of tacrolimus against *M. furfur* (Ianiri *et al*. 2017). Moreover, *M. furfur pdr10*Δ mutants did not display sensitivity to the DNA-damaging agents UV or hydroxyurea and only displayed sensitivity to the fungicide benomyl (Fig. 6C).

During BLAST searches, we noted that *M. furfur* has 2 adjacent ABC transporter-encoding genes that are orthologs of 3 adjacent ABC transporter-encoding genes in *M. sympodialis* (Fig. 6D), a *Malassezia* species that we use as a model for genomics comparison within the genus because of the high quality of its genome assembly (Zhu *et al*. 2017). Interestingly, BLASTp of these ABC transporters against *S. cerevisiae* revealed high similarity (ie E-value 0.0) with several ABC transporters, such as Pdr18, Pdr12, Pdr5, Pdr10, Pdr15, Aus1, and Pdr11. Reciprocal BLAST (BLASTp and tBLASTn) of these proteins against *M. furfur* and *M. sympodialis* finds only the aforementioned adjacent ABC transporters, which we named *Mf* (*M. furfur*) and *Ms* (*M. sympodialis*) *PDR10*_1, *PDR10*_2, and *PDR10*_3. The mutated gene in *M. furfur* corresponds to *PDR10_1*. Phylogenetic analysis revealed that ABC transporters of *M. furfur* and *M. sympodialis* cluster together in a maximum likelihood tree and are related to the *S. cerevisiae* Pdr10 ABC transporter, which is the gene designation that we selected. This analysis suggests a common duplication event of the *Malassezia PDR10* (green dot on Fig. 6E), followed by another more recent duplication in *M. sympodialis* (blue dot on Fig. 6E).

## Discussion

*A. tumefaciens*-mediated transformation is considered a “silver bullet” in functional genomics of fungi, and its main applications as well as the major discoveries that it has allowed have been recently reviewed (Idnurm *et al*. 2017a). Because *A. tumefaciens* can grow under a variety of conditions, the transformation method is versatile and has been successfully applied in a number of fungi, including those with particular nutrient requirements and that are recalcitrant to other transformation approaches, such as *Malassezia* (Ianiri *et al*. 2016; Celis *et al*. 2017).

In this report, we present the first application of forward genetics in *M. furfur*, a representative species of the fungemia-causing *Malassezia* group. The goal was to generate random insertional mutants, expose them to stress conditions to isolate those displaying sensitivity compared to the WT, and identify the corresponding T-DNA insertion sites to determine the function of the genes causing the phenotypes. Given the lack of knowledge on gene function in *Malassezia*, insertional mutants were assayed on a variety of conditions that are known to interfere with i) important cellular processes, such as those involved in plasma membrane and cell wall maintenance, growth under nutrient limiting conditions, and protein folding; ii) response to environmental stresses, such as osmotic and nitrosative stresses, UV light, elevated temperature and pH, and heavy metals; and iii) response to the antifungal FLC, which is of clinical relevance.

This loss-of-function screen allowed the characterization of 8 *M. furfur* insertional mutants (1A7, 2A8, 2F4, 6A10, 6C8, 7D9, 7F8, 7H6) that had 1 T-DNA insertion as determined by Southern blot analysis (Fig. 1) and that displayed sensitivity to one or more stress conditions (Fig. 3; Table 1). In 4 strains, the T-DNA inserted within the ORF of genes, and in another it was found to lie within a 3′ UTR (Table 1, Fig. 2 - 3), thus allowing us to define with high probability a direct link between genotype and phenotype. Clear examples of this were *M. furfur* transformants 2A8 and 1A7. Strain 2A8 was selected because of its reduced growth on minimal medium (YNB), and it was found to have a T-DNA insertion in the *ARG1* gene. Strain 1A7 was selected for its increased sensitivity to UV light, and found to have a T-DNA insertion in the *CDC55* gene. Two different approaches were employed to validate the findings of the insertional mutagenesis screen. For strain 2A8 the addition of arginine was sufficient to rescue growth to a WT level (Fig. 3A), while for strain 1A7, targeted *M. furfur cdc55*Δ mutants (Fig. 5) confirmed UV sensitivity.

In 3 other mutants of interest, the T-DNA inserted between 2 adjacent genes, and further experiments were conducted to identify the gene(s) responsible for the observed phenotype. A successful approach for strain 7H6 was gene expression analysis through RT-qPCR, which revealed downregulation of both genes flanking the T-DNA insertion (Fig. 3B). Strain 7D9 was sensitive to FLC and had a T-DNA insertion between the 3′ region of *UBC6* and 5′ region of *PDR10*, and targeted mutagenesis confirmed that the observed phenotype was due to T-DNA insertion in the promoter region of *PDR10* (Fig. 6). Lastly, the RT-qPCR approach did not allow us to define which genes was responsible for the cadmium sulfate sensitive phenotype of strain 2F4 (data not shown).

Despite the benefits of an AtMT random mutagenesis approach, analysis of the T-DNA insertion events also revealed limitations. Eleven of 19 *M. furfur* insertional mutants selected (~58%) were not useful for gene function analysis. Three transformants had 2 T-DNA insertions as determined by Southern blot analysis (Fig. 1), and although we determined at least one insertion site (Table 1), it was not possible to determine which gene was responsible for the mutant phenotype. Moreover, the insertion sites could not be identified through iPCR for 2 transformants (1F12 and 7D5). Strains 2G9 and 5D11 contained tandem T-DNA insertions, and in 4 strains (2H11, 3A1, 5F1, 5F10), chromosomal rearrangements following the integration of the T-DNA in the *M. furfur* genome were observed. Although AtMT represents a powerful method for random mutagenesis, we and other authors have commonly found non-standard T-DNA insertion events in the genome of both ascomycetous and basidiomycetous fungi [for more details see the following reviews and references within them (Michielse *et al*. 2005; Bourras *et al*. 2015; Idnurm *et al*. 2017a; Hooykaas *et al*. 2018)]. For example, in a recent study on systematic T-DNA insertion events in the red yeast *Rhodosporidium toruloides*, Coradetti and colleagues found that only 13% of mutants had regular T-DNA insertions and a total of 21% of insertions were useful to identify the genes mutated by the T-DNA (Coradetti *et al*. 2018). Moreover, in classical forward genetic screens in which loss-of-function events are selected, it is common to isolate strains with chromosomal rearrangements that originated following the insertion of the T-DNA, because these strains are generally less fit and display increased sensitivity to stress (Ianiri and Idnurm 2015). These undesirable events have been also described following AtMT of plants (Clark and Krysan 2010).

Typically in forward genetics the linkage between the T-DNA insertion and the phenotype is confirmed through: 1) sexual crosses and analysis of the phenotype in the recombinant progeny, 2) functional complementation, or 3) generation an independent targeted mutation for the gene identified (Idnurm *et al*. 2017a). Because of the lack of a known sexual cycle in *Malassezia*, and the difficulty of genetic manipulations for complementation studies, in the present study we aimed to generate *M. furfur* mutants for the genes *CDC55* and *PDR10* to validate their involvement in UV and FLC resistance, respectively. Following our previously reported protocol for targeted mutagenesis in *M. furfur* (Ianiri *et al*. 2016; Ianiri *et al*. 2017), several transformation rounds were performed, but we did not obtain any *CDC55* or *PDR10* mutants. Therefore, a system based on CRISPR/Cas9 was developed to increase homologous recombination and facilitate the generation of targeted mutants in *M. furfur*.

Because AtMT is the only effective transformation technique for *Malassezia*, a functional *CAS9* cassette and a gRNA needed to be cloned within the T-DNA of a binary vector, together with a marker for selection. For homologous recombination-mediated targeted mutagenesis, a specific gene replacement construct to serve as template to repair the BSB was also necessary. Cloning of all the required components within the T-DNA of one binary vector is technically challenging and time consuming. In one study, Kujoth and colleagues generated a large T-DNA that included one or more gRNA, a Cas9 expression cassette, and a gene marker, and successfully applied this system in gene editing strategies through NHEJ in *Blastomyces dermatitidis* (Kujoth *et al*. 2018). For CRISPR/Cas9 in *Leptosphaeria maculans*, Idnurm and colleagues reported a system based on 2 binary vectors, one with *CAS9* and a marker, and the other with gRNA and another marker, that could be successfully delivered at the same time through co-transformation employing *A. tumefaciens* to perform efficient gene editing through NHEJ (Idnurm *et al*. 2017b). For CRISPR/Cas9 of *Malassezia*, we opted for a system that would be suitable for targeted gene replacement through homologous recombination based on co-transformation of *M. furfur* with 2 *A. tumefaciens* strains, one bearing the binary vector with the HDR gene deletion allele, and another with a binary vector engineered for the CRISPR/Cas9 system without a gene marker. The rationale for generating a marker-free binary vector was to: 1) have a CRISPR/Cas9 transient expression system with a reduced rate of *CAS9* and/or gRNA ectopic integration in the host genome, similar to a system developed for *C. albicans* and *C. neoformans* (Min *et al*. 2016; Fan and Lin 2018); 2) allow further genetic manipulation of the *NAT*-generated mutant using the other *Malassezia*-specific gene marker available, which encodes for resistance to neomycin; and 3) reduce recombination within the actin promoter and terminator regions of the *NAT* and *NEO* gene markers.

Considering gRNA expression, the choice of an appropriate promoter has represented a major challenge for the application of CRISPR/Cas9 technology in fungi. Currently, common approaches include the use of a strong promoter recognized by RNA polymerase II, such as that of the *ACT1* or *GDP1* genes, coupled with a hammerhead ribozyme and/or hepatitis delta virus ribozyme for gRNA excision (Idnurm *et al*. 2017b; Kujoth *et al*. 2018); the use of RNA polymerase III promoters, such as the U6 promoters of small nuclear RNA used for *C. neoformans* (Wang *et al*. 2016; Fan and Lin 2018); or the promoters of the tRNA or rRNA with or without ribozymes (Shi *et al*. 2017). In this study, we first tested a strategy based on the use of the 5S rRNA promoter of *M. sympodialis* (Fig. 4B). While we were working on developing this system, Zheng and colleagues reported a similar approach in *A. niger*, demonstrating high efficiency of gene editing using both the 5S rRNA promoter alone or combined with the HDV ribozyme (Zheng *et al*. 2018). Cas9 expression is usually achieved using a strong promoter and terminator; in this study the regulatory regions of the histone *H3* gene of *M. sympodialis* served this purpose (Fig. 4A). During the first CRISPR/Cas9 attempt, we were able to generate *M. furfur cdc55*Δ mutants. Surprisingly, both molecular and phenotypic analysis revealed a homologous recombination rate of 98% (Fig 4C, and Fig. S1). This high rate of homologous recombination is similar to that in the study of Zheng and colleagues (Zheng *et al*. 2018) and other CRISPR/Cas9-mediated gene deletion approaches (Fan and Lin 2018). Given these positive results, the use of ribozymes flanking the 5S rRNA promoter was not tested. Phenotypic analysis confirmed the involvement of *CDC55* in UV resistance, and further assays revealed sensitivity of the *M. furfur cdc55*Δ mutant to benomyl and hydroxyurea (Fig. 5D), which also induced an abnormal bud morphology (Fig. 5E). This indicates a conserved function of the cell division cycle protein Cdc55 in *M. furfur* and *S. cerevisiae*.

This CRISPR/Cas9 technology was then tested for targeted mutagenesis of another gene of interest, *PDR10*. Corroborating results obtained for *CDC55*, we were able to promptly obtain *pdr10*Δ mutants, although the rate of homologous recombination was lower for this gene. This could be due to several factors, such as shorter flanking regions of ~800 bp used in the HDR *pdr10*Δ*::NAT* template, the length of the *PDR10* gene (more than 4 Kb), the genomic location, or to lower activity of the *PDR10*-specific gRNA. Analysis of *M. furfur pdr10*Δ mutants revealed an unexpected specificity of *M. furfur PDR10* for resistance to the clinical-relevant drug FLC, and to the antifungal agent benomyl (Fig. 6C). While there are multiple studies on the pleiotropic drug resistance function of ABC transporters in non-pathogenic (*S. cerevisiae*) and pathogenic (*C. albicans*) yeasts (Sipos and Kuchler 2006; Coste *et al*. 2008; Paul and Moye-Rowley 2014), in these cases specific analysis of Pdr10 in response to several drugs has yet to be performed, and therefore it is not possible to provide a detailed comparison analysis that supports conserved or divergent functions of *PDR10* in *M. furfur*. A recent study reported the involvement of *S. cerevisiae* Pdr10 in double-strand break repair via sister chromatid exchange (MuÑoz-GalvÁN *et al*. 2013), but we could not confirm this function in *M. furfur* because of the lack of sensitivity of *pdr10*Δ mutants to DNA-damaging agents (Fig. 5C). Further bioinformatics analyses revealed that Pdr10 is the only ABC transporter present in the genome of two *Malassezia* species, and that it underwent ancestral and more recent gene duplication events (Fig. 6D-E). This suggests profound differences with other fungi, and further studies are needed to elucidate the evolution and specific roles of these *Malassezia* ABC transporters in resistance to chemicals and their network of interactions.

Our understanding of *Malassezia* genetics is still limited, and the T-DNA-mediated random insertional mutagenesis applied in this study coupled with a novel and efficient CRISPR/Cas9 system represent straightforward approaches to advance molecular genetics in this understudied organism. Indeed, while T-DNA-mediated random insertional mutagenesis is of particular relevance to discover novel gene functions, such as the UV sensitive phenotype of *CDC55*, or the FLC sensitivity due to mutations in the genes *SIP5* and *ADY2*, the efficiency of CRISPR/Cas9 is a critical requirement to perform large-scale analyses while also validating the results of the genetics screen.

Historically *Malassezia* research has been hampered by the fastidious nature and particular growth requirements of species within this genus and by their difficult identification and classification. Nevertheless, in addition to the available genome sequence and annotation of most *Malassezia* species, the recent introduction of animal models to study *Malassezia* interactions with the skin and the gastrointestinal tract (Limon *et al*. 2019; Sparber *et al*. 2019), and the development of this novel CRISPR/Cas9 system and other existing molecular technologies (Ianiri *et al*. 2016; Celis *et al*. 2017; Ianiri *et al*. 2017) represent key scientific advances to study the biology and pathophysiology of *Malassezia*, the main fungal inhabitants of mammalian skin.

## ACKNOWLEDGMENTS

We thank Sheng Sun for assistance with Southern blot analysis, and Tom Dawson for sharing unpublished genome and RNAseq information prior to publication. We thank Ci Fu, Shelby Priest and Cecelia Wall for critical comments on the manuscript.

---

**Fig. S1.**
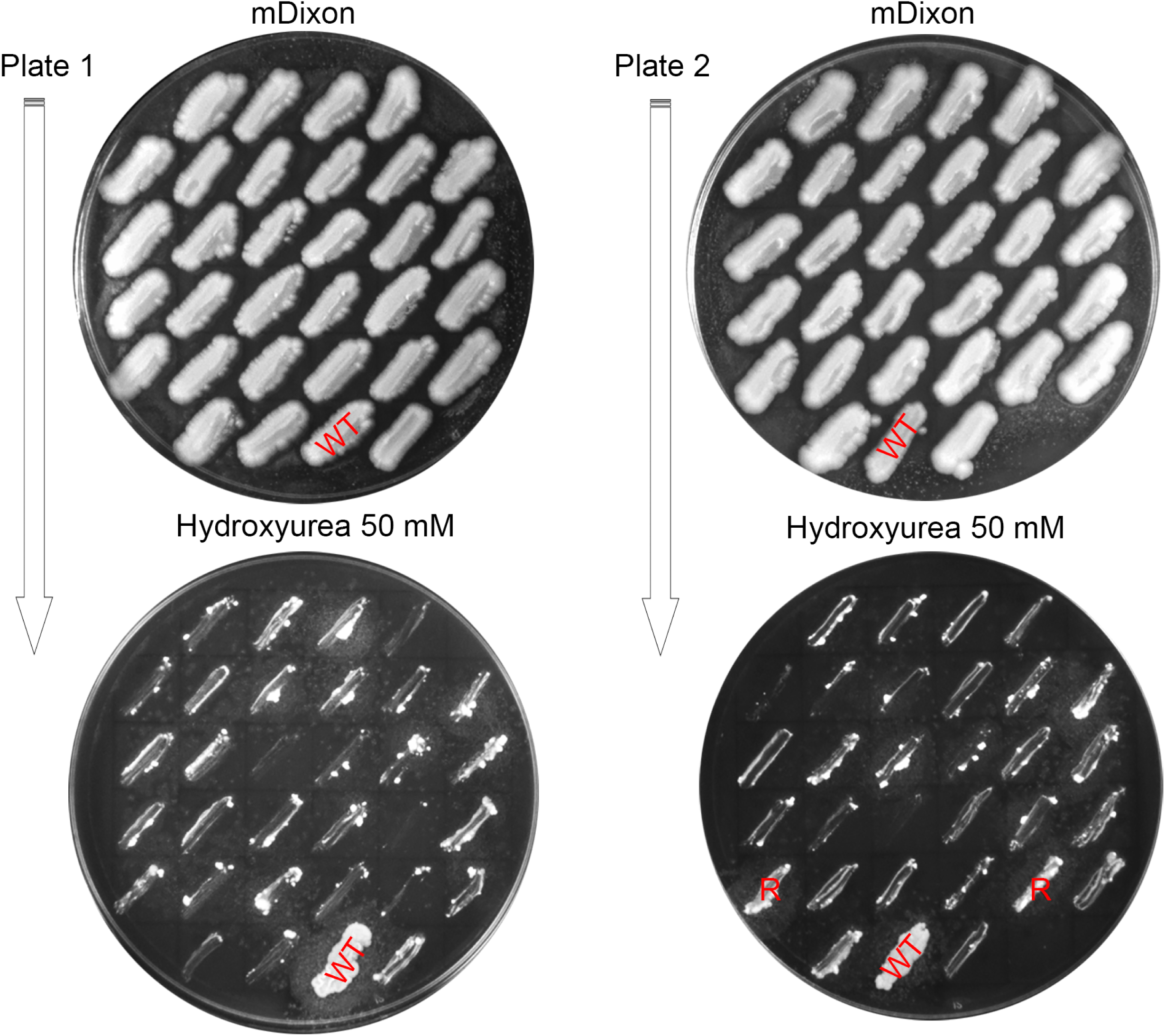
Phenotypic characterization of putative *M. furfur* cdc55Δ mutants on mDixon + hydroxyurea (50 mM). The *M. furfur* WT strain and 2 transformants that showed increased resistance (R) to hydroxyurea are indicated.

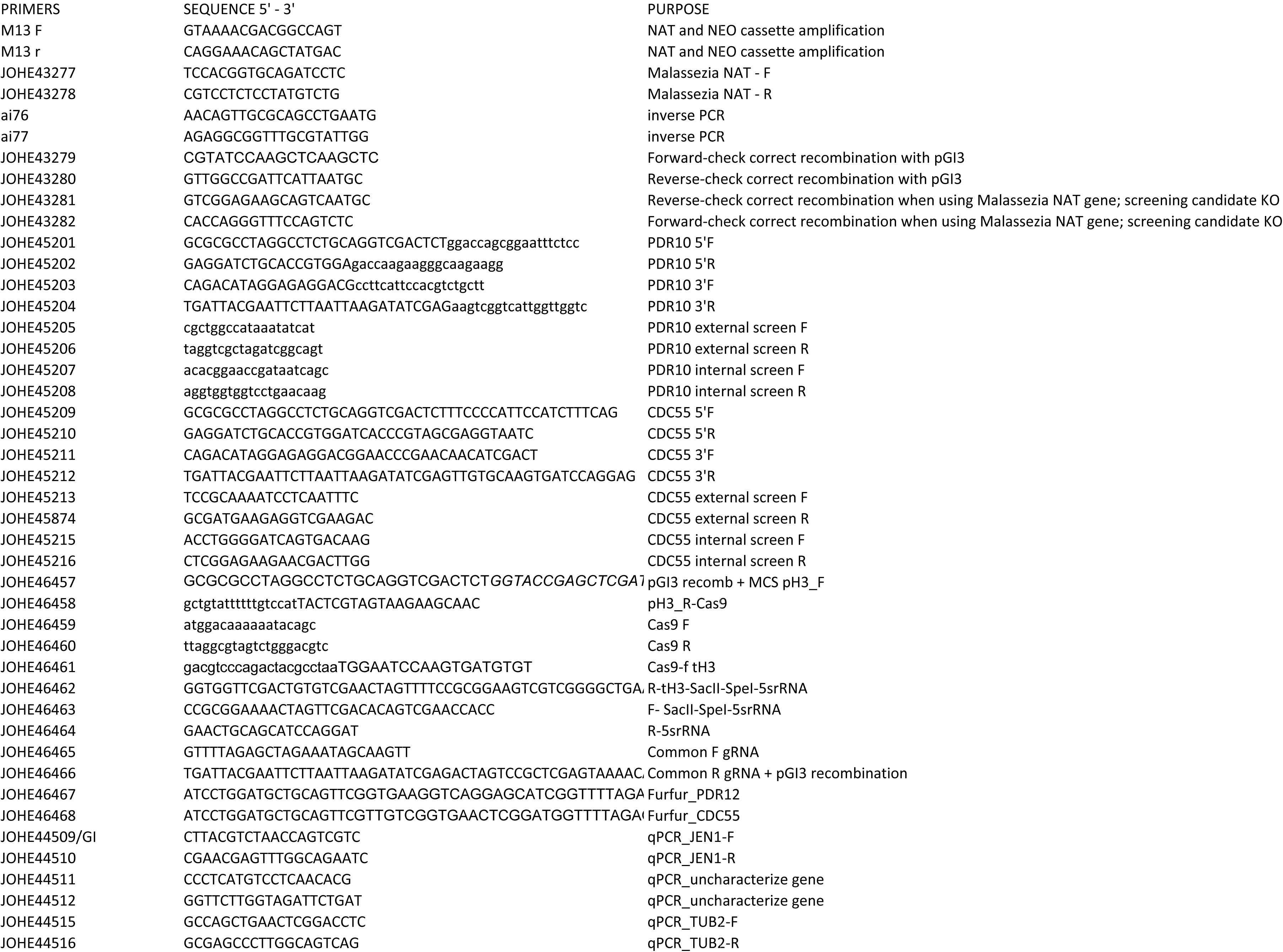

